# GuPPy, a Python toolbox for the analysis of fiber photometry data

**DOI:** 10.1101/2021.07.15.452555

**Authors:** Venus N. Sherathiya, Michael D. Schaid, Jillian L. Seiler, Gabriela C. Lopez, Talia N. Lerner

## Abstract

Fiber photometry (FP) is an adaptable method for recording *in vivo* neural activity in freely behaving animals. It has become a popular tool in neuroscience due to its ease of use, low cost, the ability to combine FP with freely moving behavior, among other advantages. However, analysis of FP data can be a challenge for new users, especially those with a limited programming background. Here, we present <u>**Gu**</u>ided <u>**P**</u>hotometry Analysis in <u>**Py**</u>thon (GuPPy), a free and open-source FP analysis tool. GuPPy is provided as a Jupyter notebook, a well-commented interactive development environment (IDE) designed to operate across platforms. GuPPy presents the user with a set of graphic user interfaces (GUIs) to load data and provide input parameters. Graphs produced by GuPPy can be exported into various image formats for integration into scientific figures. As an open-source tool, GuPPy can be modified by users with knowledge of Python to fit their specific needs.

## INTRODUCTION

Fiber photometry (FP) is an adaptable method for recording *in vivo* neural activity in freely behaving animals. FP is a relatively new technique, with first reports published in the last decade ^1–3^, which takes advantage of concurrent advances in fluorescent reporters for calcium and other biomolecules of interest ^4^. Unlike traditional imaging of fluorescent reporters, FP does not create a spatially resolved image. Rather, photons are collected through a single fiber optic brain implant, without information as to their origin. Therefore, FP provides a bulk measurement of reporter activity near the fiber optic probe but cannot offer single cell resolution. Despite this limitation, FP has grown in popularity amongst neuroscientists due to advantages compared to other methods. Among these advantages are the ease of use, low cost, the ability to combine FP with freely moving behavior, the ability to image in two colors, and the ability to image from multiple brain sites simultaneously.

FP requires a single stereotaxic surgery in which a fiber optic probe is implanted in a brain region of interest. Light can be delivered through the probe to excite fluorescent reporters (expressed via viral methods or in transgenic animals), and light emitted from the reporters can be collected back through the same probe. The head apparatus size is much smaller compared to other *in vivo* methods such as microendoscopes (miniscopes) or electrophysiological recordings that require large headcaps. Wireless adaptions of FP technology are also in development ^5^. As a result, FP is versatile in its application and compatible with a variety of behavioral paradigms. Because it is light-based, recordings are not susceptible to electrical artifacts. Tissue damage to surrounding structures is also reduced as compared to other methods: fiber optic probes are generally 200-400um in diameter as compared to 500um-2mm GRIN lens implants for imaging. For measurements of activity from deep brain structures, these diameter differences are significant, resulting in much less damage to overlying brain structures. The small probes required for FP also allow for multiple implants per animal and, thus, multi-site recordings from a single animal are possible, with up to 7 having been demonstrated in the literature ^6,7^. Tapered and submicrometer probes have been used for multi-site recordings as well ^8,9^.

The adoption of FP in labs around the world has been facilitated by an increasing number of affordable “off-the-shelf” commercial options for hardware that require minimal assembly or alignment of optical parts. Companies such as Tucker Davis Technologies and Doric Lenses were among the first to market, and the list of options is growing. The ease of data collection in FP has led to the expansion of its use, but the pipelines for data analysis are much less standardized. In the existing literature using FP, data analysis between groups has varied and has mostly been reported as “custom analysis.” To date, only one published analysis tool box exists ^10^. Although very effective, this tool limits users to MATLAB software run on the Windows operating system. There remains a demand for consistent, user-friendly, and low-cost analysis tools. Here we present <u>**Gu**</u>ided <u>**P**</u>hotometry Analysis in <u>**Py**</u>thon (GuPPy), a free and open-source tool designed to guide users with minimal programming knowledge in the analysis of FP data. GuPPy is provided as a Jupyter notebook, a well-commented interactive development environment (IDE) designed to operate across platforms. GuPPy provides users with limited programming knowledge a well-documented workflow, while allowing those with greater programming knowledge the ability to flexibly adjust all their analysis parameters as desired. Here, we describe GuPPy’s function and capabilities as well as plans for future development. Access to GuPPy is freely provided for immediate use, with the goal of standardizing FP analysis and aiding in the adoption of FP for new users.

## RESULTS

### General Principles

In developing GuPPy, we sought to provide a tool for FP data analysis that is based on a free platform (Python) and that can be used across the Windows, Mac, and Linux operating systems. In designing GuPPy, we kept the following goals in mind:

- To create a flexible tool for the analysis of FP data recorded during many different types of behavioral experiments
- To create a tool that guides users unfamiliar with Python through the analysis
- To create an open-source tool that experienced programmers can adapt for their own specific needs, avoiding unnecessary duplications of effort by programmers in different labs
- To allow users to produce attractive graphical outputs for scientific figures

GuPPy was developed to analyze FP data recorded using Tucker Davis Technologies (TDT) processors and Synapse software (see Methods), although future versions may incorporate additional options for input from other systems. Herein, we give a high-level description of the analysis currently possible in GuPPy without modification of the underlying code. To access GuPPy and to view a detailed, step-by-step user guide, please visit our <u>Github Wiki page</u> (https://github.com/LernerLab/GuPPy/wiki).

### Loading Data and Setting Input Parameters

FP datasets recorded using TDT systems are stored in tanks and blocks. A tank is generally used to hold the data from multiple recording sessions associated with a particular experiment. Blocks are unique folders within tank directories. Within a block, different stores can record FP signals and behavioral event timestamps. Although TDT stores are the best option for recording behavioral timestamps that are properly synchronized with FP data recorded by a TDT processor, in the case that behavioral timestamps are recorded elsewhere, GuPPy can also accept event timestamps in a csv file format.

GuPPy supports the analysis of a single FP recording session (single processing) and the simultaneous analysis of multiple recording sessions (batch processing). Within GuPPy, an ‘Input Parameters’ GUI is provided, which allows the user to select the folder(s) where recorded TDT files are stored. Selection of a single folder (a.k.a. block) will invoke single processing and selection of multiple folders will invoke batch processing. To use a csv file to input event timestamps, the file simply needs to be stored in the same folder as the associated TDT files. GuPPy will automatically look for any csv event timestamp files while reading the data.

The Input Parameters GUI also asks the user to set a variety of parameters that will be used in the analysis. The user can specify various types of artifact correction to execute, including removing parts of the data containing artifacts and using an isosbestic control wavelength to normalize data (these processes are described in more detail in the next section). The user can also specify parameters to be used for analysis, for example to create peri-stimulus/event time histograms (PSTHs).

### Specifying Storenames

After saving the input parameters, the user must specify the storenames to analyze using the ‘Open Storenames’ GUI. When the user opens this GUI, single or multiple browser windows will pop up based on whether the user is performing single processing or batch processing. Instructions are provided by GuPPy on how to specify storenames for analysis (i.e. those stores within the TDT blocks containing the FP data). GuPPy will extract the data from the stores containing the isosbestic control and signal channels. After extraction of the data, a linear digital filter is applied backwards and forwards to both the channels to reduce high frequency noise without time-shifting. The resulting traces for the control and signal channels will be displayed for the user to inspect before proceeding with analysis.

### Data Analysis

#### Artifact Removal

Once the data is loaded, the analysis workflow begins with artifact correction. First, a set amount of time (specified as an input parameter, e.g. 1s) can be removed from the beginning of the recording. This feature is useful if, for example, the light sources for exciting fluorescent reporters turn on at the beginning of the recording and create an artifact (Fig. 1a). Second, an option to ‘Remove Artifacts’ allows users to select portions of their recordings to include and exclude from in the analysis. This feature is useful if large artifacts occurred in the middle of a recording (Fig. 1b). If these artifacts are not dealt with, it will be difficult to properly fit the control and signal channels. To remove artifacts, the user manually selects the start and end points for all the chunks of data they desire to keep. The coordinates for selected portions of the data will be saved to use in the next analysis steps. If only a limited amount of time within a recording is disrupted by such an artifact, the recording can in many cases be salvaged by this mechanism.

**Figure 1.**
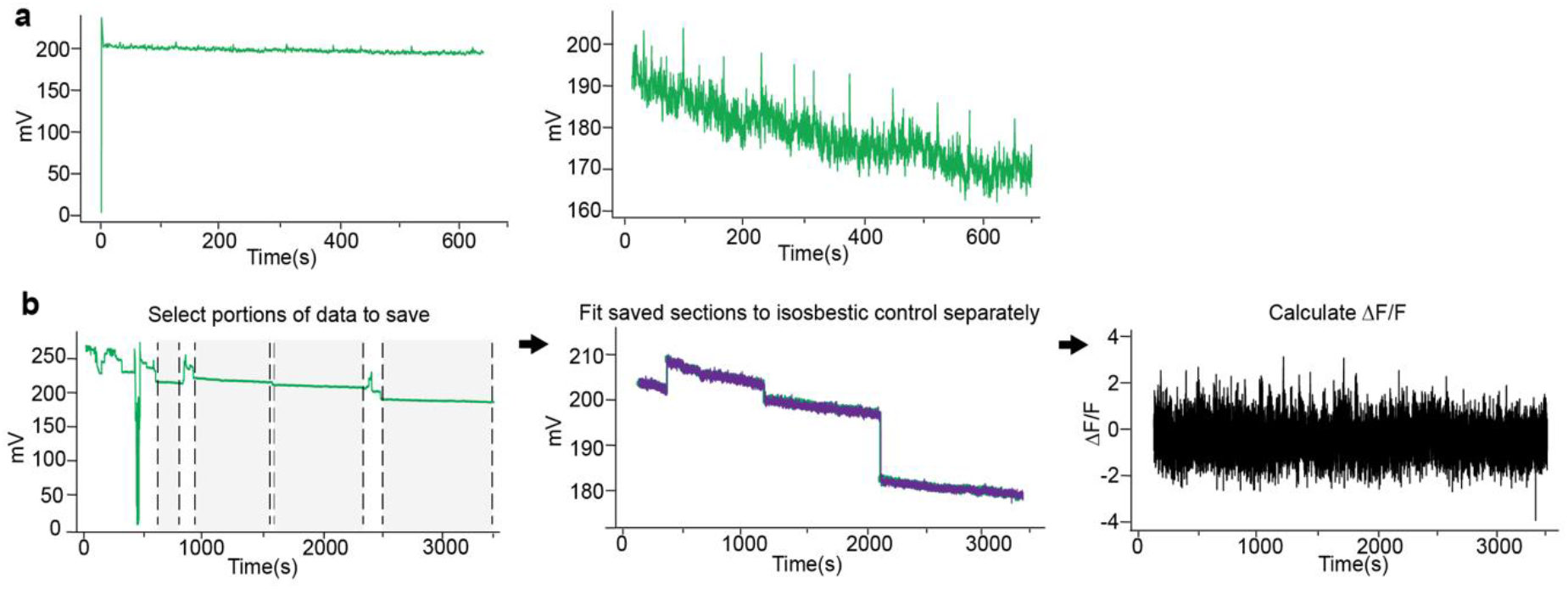
Artifact Correction Options in GuPPy. **a)** A set amount of time at the beginning of a recording can be removed. This artifact correction option provides a fast and easy way to remove an artifact regularly occurring when recordings initiated. An example of such an artifact, which occurs when light sources turn on automatically at the beginning of a recording, is shown on the left. Clean data after the removal of this artifact (the first 1s of the recording) is shown on the right. **b)** GuPPy allows user-selected artifact correction to remove artifacts that can sometimes occur unpredictably mid-recording. An example of a recording where this problem occurred is shown. To salvage much of the data from such a recording, the user can manually select portions to keep, and GuPPy will separately fit the control and signal channels for each section and then concatenate the saved sections for further processing, preserving behavioral timestamp alignments.

#### Generating ΔF/F and z-scores

GuPPy is programmed to use a control isosbestic wavelength recorded by users to remove smaller movement artifacts as well as bleaching artifacts when calculating the ΔF/F (as described in Lerner et al, 2015; Fig. 2). The isosbestic wavelength includes artifacts, but not calcium-dependent events (in the case of a calcium sensor), and so can be subtracted out from the signal and used to normalize the data to determine changes from baseline fluorescence (ΔF/F). In the Input Parameters GUI, we have included the option to display data as ΔF/F or as a deviation of the ΔF/F signal from its mean (z-score). Z-scores are useful for combining data across multiple mice or sessions.

**Figure 2.**
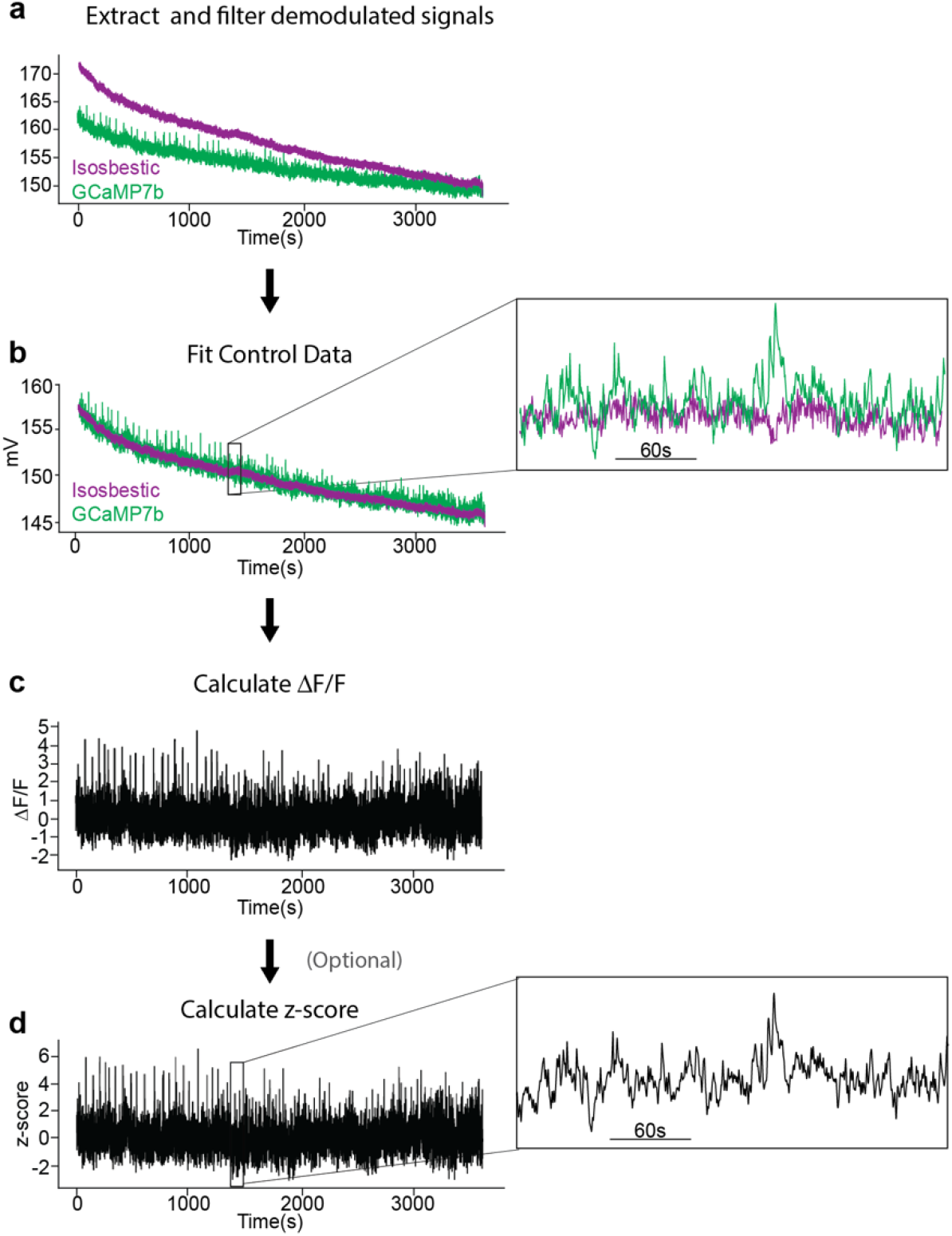
Raw data extraction, fitting and analysis. **a)** Example data from a single 1-hour recording. Calcium in dopamine axons in the dorsomedial striatum was recorded using GCaMP7b. An isosbestic (ex405nm-purple) and calcium-dependent (ex465nm-green) signal were recorded simultaneously. **b)** After both data streams are filtered and demodulated, the isosbestic control data is fit to the signal. The fitted control is then used to calculate **c)** change in fluorescence (ΔF/F) and **d)** z-score.

A fitted control channel is obtained by fitting the control channel to signal channel using a least squares polynomial fit of degree 1. ΔF/F is computed by subtracting the fitted control channel from the signal channel, and dividing by the fitted control channel:

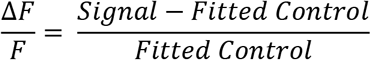

If the z-score option was chosen by the user, a z-score signal is then computed using the formula:

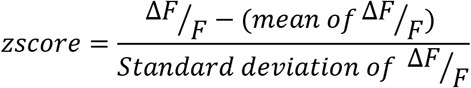

If the artifacts removal functionality described in the previous section was used to exclude portions of a recording, ΔF/F will be calculated separately for each individual chunk of saved data, and then these chunks will be combined to compute the final z-score of the whole trace. During the process of cutting and pasting chucks together, associated behavioral event timestamps are also aligned to make sure later computations do not misalign FP data with the behavioral timestamps.

#### Average Amplitudes and Frequencies of Events Within Entire Recording Sessions

The average amplitude and frequency of transients in the z-score trace computed for the whole session can be calculated by GuPPy. GuPPy uses a user-defined moving window (default is 15s) for thresholding transients. High amplitude events – defined as events with amplitudes greater than two times the median absolute deviation (MAD) above the median of the moving window – are filtered out and the median of the resultant trace is calculated. Peaks with a local maxima greater than three MADs of the resultant trace are counted as transients. While somewhat arbitrary, these criteria generally select peaks that align with a human observer’s judgement for detecting transients (Fig. 3). This approach for transient detection was adapted from analyses performed in recent publications ^11,12^.

**Figure 3.**
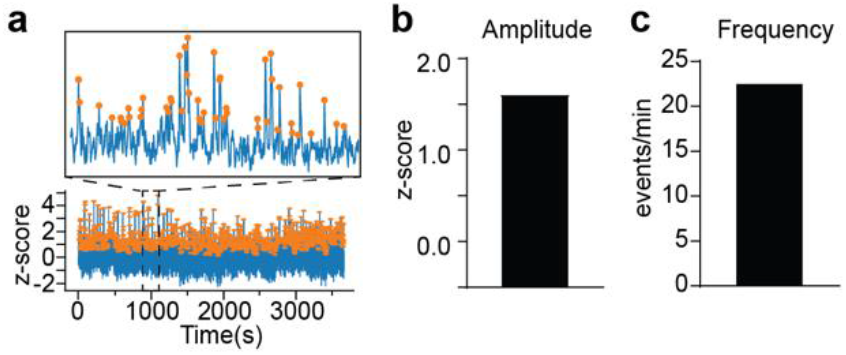
Peak detection. **a)** A trace demonstrating z-score peak detection during a 30 minute recording. Orange dots represent peaks detected by the software. The inset shows an expanded portion of the data. During peak detection, **b)** average peak amplitude and **c)** average peak frequency are calculated and these values are returned to the user for analysis.

#### PSTHs

PSTHs are an efficient way to examine responses to discrete behavioral events over repeated trials. GuPPy will compute PSTHs based on a user defined window [A,B] set around each event timestamp. The first parameter, A, represents time (in seconds) before the event timestamp and the second parameter, B, represents time (in seconds) after the event timestamp to be used when calculating the PSTH. A baseline correction can be performed within the PSTH calculation if the user selects this option. If baseline correction is selected, then GuPPy takes the mean value in a user-defined baseline window for each trial within the PSTH and subtracts it from that trial’s trace before averaging the traces together to display the final PSTH.

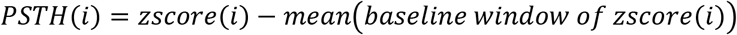

where *i* represents each event included in the PSTH

The average of all trials in the PSTH vector is calculated after baseline correction. The user can then calculate the area under the curve (AUC) and the peak of this PSTH within a defined window. AUC is computed using the trapezoidal method, beginning from the event timestamp (at time 0 of the PSTH) and continuing until the average value of the PSTH vector goes below baseline and remains below the baseline for 0.5 seconds.

#### PSTH Visualization

PSTHs can be visualized by GuPPy in various ways. The user can browse through individual events making up the PSTH and display the mean PSTH with or without the individual traces shown (Fig. 4a–b). The user can select to plot the PSTH for one event, or to plot PSTHs for multiple events together on the same graph (Fig. 4c).

**Figure 4.**
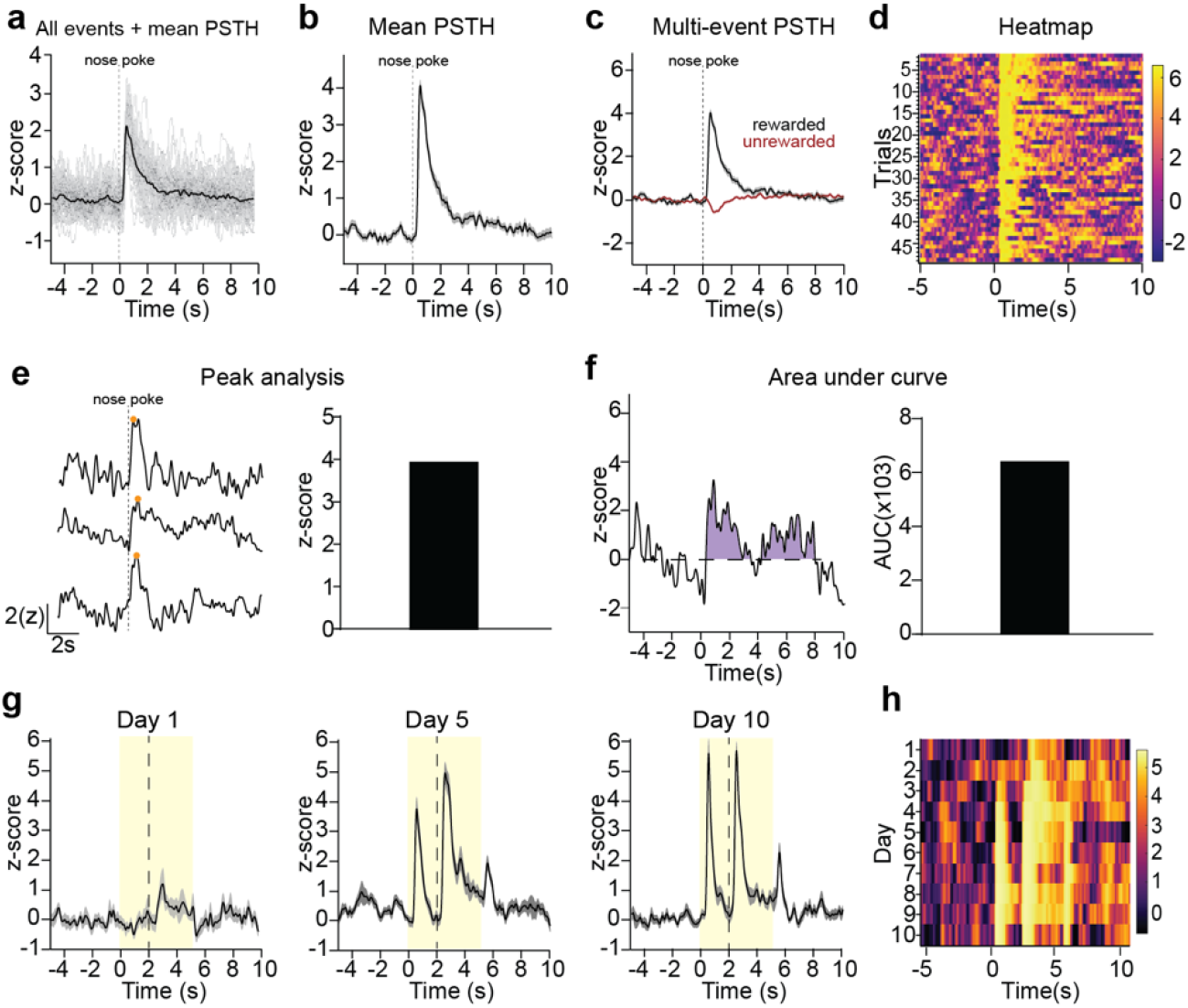
Visualization of data from individual recordings. **a-f)** A representative recording of GCaMP7b-expressing dopamine axons in dorsomedial striatum during an instrumental behavioral task in which animals nosepoke for reward is used to illustrate data visualization and analysis options. **a)** Peri-stimulus/event time histogram (PSTH) for rewarded nosepokes, in which the mean z-score (black) is plotted along with all individual trials (gray). **b)** Mean PSTH for rewarded nosepokes plotting mean z-score with standard error. **c)** A multi-event PSTH plot comparing average responses to rewarded (black) vs unrewarded (red) nosepokes. **d)** Heatmap of all individual trials for rewarded nosepokes. Trials are shown in chronologically order from top to down. The color scheme for heatmaps in GuPPy is easily modifiable by the user. **e)** Example traces illustrating peak detection. Orange dots indicate the detected peaks (left) and an average peak amplitude for all rewarded nosepoke trials is calculated (right). **f)** An example trace illustrating area under curve (AUC) calculations. The AUC counted is colored in purple (left) and the average AUC for all rewarded nosepoke trials is calculated (right). **g-h)** A representative recording of dLight1.3b in nucleus accumbens during a Pavlovian conditioning task is used to illustrate additional data visualization options comparing recordings across days. **g)** PSTHs for days 1, 5, and 10 of Pavlovian training are show separately. A cue (yellow box) predicted reward (dotted line). **h)** A heatmap is used to display average PSTHs across days 1-10 in a compact format, allowing easy visualization of the emergence of cue-evoked dopamine with training.

GuPPy can also produce heatmaps to display the individual trials making up a PSTH in a compact format (Fig. 4d). The color spectrum used for the heatmap can be adjusted by the user. Displaying individual trials in this manner is useful for visualizing whether the average PSTH trace produced from a recording session arises from a consistent response, or from some other pattern such as a progression of responses through the training session. For example, we show here the responses of dopamine axons to a rewarded nosepoke as a heatmap showing all trials, demonstrating that this response is seen consistently throughout all trials in the recording session for a well-trained mouse (Fig. 4d).

GuPPy can also produce heatmaps that display average PSTHs traces from different recording sessions, e.g. to display changes in the patterns of activity seen across days. For example, here we show a mouse being trained to associate a Pavlovian cue (lever insertion) with a reward delivery (sucrose pellet; Fig. 4g). The heatmap across days of training allows one to easily visualize the emergence of cue-evoked dopamine in a much more compact manner than displaying PSTHs for each day separately (Fig. 4h).

#### Group Analysis

One common desire by users is to average together results from multiple animals, or multiple recording sessions to determine group differences (Fig. 5). This type of “group analysis” is therefore an important feature, which GuPPy offers. After the analysis of individual sessions has been run, the user can select all the folders of data they want to average together in the Input Parameters GUI and set the ‘Average Group’ parameter to ‘True’. Upon running the data analysis again, the average results will be computed. Visualization of these average results is possible by setting the ‘Visualize Average Results’ parameter to ‘True’.

**Figure 5.**
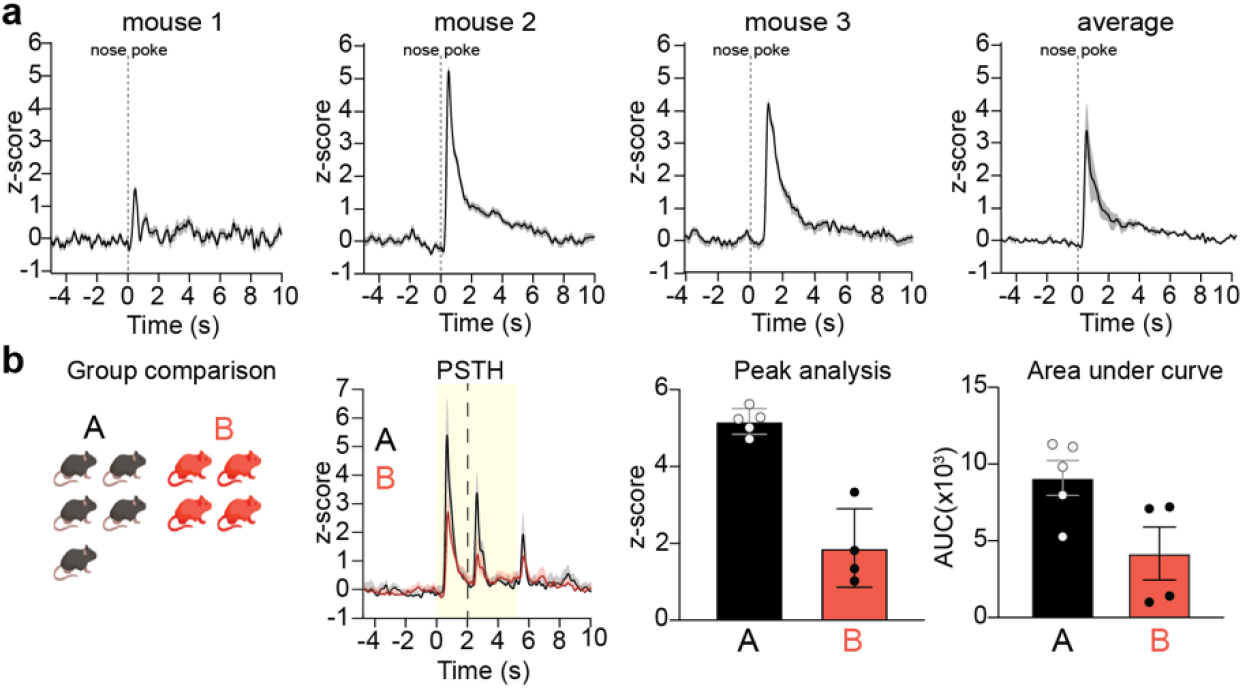
Visualization of data from group analyses. GuPPy allows users to compile data from multiple mice according to their assigned experimental groups. **a)** Three individual representative recordings and the average response from all 3 mice are shown. Recordings are from GCaMP7b-expressing dopamine axons in dorsomedial striatum during an instrumental behavioral task in which mice nosepoke for reward (same as data from Fig. 4a–f). **b)** Representative group analysis from 2 groups of mice (A-black and B-red). Recordings are from dLight1.3b expressed in nucleus accumbens (same as data from Fig. 4g–h). Mice were trained on a Pavlovian conditioning task, in which a cue (yellow box) predicted reward (dotted line). Data is plotted as PTSH, peak z-score, and AUC.

GuPPy’s ability to plot multiple PSTHs (as in Fig. 4c) also lets the user plot PSTHs for the same event in different groups of animals on the same graph, which can be useful for group analyses (Fig. 5b).

### Outputs

All the results at each point in the analysis are saved in either hdf5, csv or h5 format. Reading and writing data into hdf5 or h5 format is faster and consumes less memory. Saving in these formats also has the advantage of making these files accessible in other programming languages, including MATLAB.

All the data visualizations created by GuPPy can be saved as PNGs, SVGs, or both. PNG is a raster graphics format that allows exporting high-resolution images that can be displayed by many programs such as Microsoft Word and basic photo viewers. SVG is a vector graphics format that allows images to be edited in Adobe Illustrator, Inkscape, Microsoft Powerpoint, or other graphics software for flexible incorporation into larger scientific figures.

## DISCUSSION

The growing popularity of FP has created a need for accessible analysis tools. We were motivated to develop GuPPy to standardize our own lab’s analyses, to allow new users to engage quickly with the technology, and to enable sharing across the community of FP users to avoid unnecessary duplications of effort. Currently, only one other FP analysis tool, pMAT ^10^, is publicly available. While valuable, this tool is programmed in MATLAB, a platform that requires a paid subscription to use, and works only with a Windows operating system. In developing GuPPy, we sought to provide an additional tool that is based on a free platform (Python) and that can be used across the Windows, Mac, and Linux operating systems. A user can interact with GuPPy through GUIs that do not require a working knowledge of Python. Users with knowledge of Python are free to make improvements and adjustments to this open-source tool.

As with any software tool, continued development of GuPPy can improve its functionality. In addition to providing open-source code to the community, we are planning continued internal development. We plan to release new updates following this initial launch and welcome feedback or suggestions. To report bugs or to request new features in the tool, please raise an issue on <u>GuPPy GitHub</u> page or join the <u>Gitter</u> chat room to ask questions.

## MATERIALS AND METHODS

### Mice

Male and female *WT* (C57BL/6J) and (DAT)::IRES-Cre knockin mice (JAX006660) were obtained from The Jackson Laboratory and crossed in house. Only heterozygote transgenic mice, obtained by backcrossing to C57BL/6J wildtypes, were used for experiments. Adult mice at least 10 weeks of age were used in all experiments. Mice were group housed under a conventional 12h light cycle (dark from 7:00pm to 7:00am) with *ad libitum* access to food and water prior to behavioral training. During training, mice were food restricted to 85% of their *ad libitum* weight. All experiments were approved by the Northwestern University Institutional Animal Care and Use Committee. All methods were carried out in accordance with relevant guidelines and regulations. Insofar as they are relevant, the methods reported below are in accordance with ARRIVE guidelines (however, since only example data is shown to illustrate software functionality, many study design and statistical details are not applicable).

### Stereotaxic Surgery

Viral infusions and optic fiber implant surgeries took place under isoflurane anesthesia (Henry Schein). Mice were anesthetized in an isoflurane induction chamber at 3-4% isoflurane, and then injected with buprenorphine SR (Zoopharm, 0.5 mg/kg s.q.) and carpofen (Zoetis, 5 mg/kg s.q.) prior to the start of surgery. Mice were placed on a stereotaxic frame (Stoetling) and hair was removed from the scalp using Nair. The skin was cleaned with alcohol and a povidone-iodine solution prior to incision. The scalp was opened using a sterile scalpel and holes were drilled in the skull at the appropriate stereotaxic coordinates. Viruses were infused at 100 nl/min through a blunt 33-gauge injection needle using a syringe pump (World Precision Instruments). The needle was left in place for 5 min following the end of the injection, then slowly retracted to avoid leakage up the injection tract. Implants were secured to the skull with Metabond (Parkell) and Flow-it ALC blue light-curing dental epoxy (Pentron). After surgery, mice were allowed to recover until ambulatory on a heated pad, then returned to their homecage with moistened chow or DietGel available. Mice then recovered for three weeks before behavioral experiments and fiber photometry recordings began.

### Fiber Photometry

Fiber photometry data used here to illustrate the functionality of GuPPy was collected using a fiber photometry rig with optical components from Doric Lenses controlled by a real-time processor from Tucker Davis Technologies (TDT; RZ5P). TDT Synapse software was used for data acquisition. 465nm and 405nm LEDs were modulated at 211 or 230 Hz and 330 Hz, respectively. LED currents were adjusted in order to return a voltage between 150-200mV for each signal, were offset by 5 mA, and were demodulated using a 4 Hz low-pass frequency filter. Behavioral timestamps were fed into the TDT processor as TTL signals from the operant chambers (MED Associates) for alignment with the neural data.

For experiments shown in Fig. 4A–F and Fig. 5, mice received infusions of 1μl of AAV5-CAG-FLEX-jGCaMP7b-WPRE into medial SNc (AP −3.1, ML 0.8, DV −4.7) and a fiber optic implant was placed in DMS (AP 0.8, ML 1.5, DV −2.8). The data shown is from a recording session during random interval (RI60) training.

For experiments shown in Fig. 4G–H, mice received infusions of 500nl of AAV9-CAG-dLight1.3b virus into NAc core (AP 1.5, ML 0.9, DV −4.1) and a fiber optic implant was placed in NAc core at the same coordinates. The mouse was trained on a Pavlovian task in which the mouse was trained to associate a 5 sec lever cue with the delivery of a sucrose pellet. The sucrose pellet was delivered 2 sec after the initiation of the lever cue.

### GuPPy Code and Tutorials

GuPPy code is posted on Zenodo (https://zenodo.org/badge/latestdoi/382176345) and on GitHub (https://github.com/LernerLab/GuPPy). Any future updates to GuPPy will be available at these sources. Along with the code, a detailed step-by-step user guide is provided on GitHub (https://github.com/LernerLab/GuPPy/wiki). GuPPy was developed using Python 3.6 but it will work for Python 3.7 or 3.8 as well.

### Computer System Requirements

GuPPy can be run on any operating system (Windows, Mac, Linux). Users must download Anaconda compatible with their operating system (https://www.anaconda.com/products/individual#macos). There are no special computer system requirements, however, we recommend at least 8GB RAM and a 2-4 core processor to allow fast batch processing for larger datasets e.g. batch processing more than 10 recording sessions.

## ACKNOWLEDGMENTS

We thank all the members of the Lerner Laboratory for helpful discussions and critical feedback on the development of this analysis tool. We thank David Barker, M. Gunes Kutlu, Erin Calipari, Vaibhav Konanur, and Mitch Roitman for sharing data used to test GuPPy. This work was supported by an NIH New Innovator Award (DP2MH122401) to T.N.L and by a NIDA Specialized Center Grant (P50DA044121) supporting G.C.L.

## AUTHOR CONTRIBUTIONS

**VNS:** Conceptualization, Methodology, Software, Formal Analysis, Writing – Original Draft, Visualization

**MDS:** Methodology, Writing – Original Draft, Visualization

**JLS:** Methodology, Investigation, Writing – Review & Editing

**GCL:** Investigation

**TNL:** Conceptualization, Resources, Writing – Review & Editing, Supervision, Project administration, Funding Acquisition

## REFERENCES

1. Cui, G. et al. Concurrent activation of striatal direct and indirect pathways during action initiation. Nature 494, 238–242 (2013).

2. Gunaydin, L. A. et al. Natural Neural Projection Dynamics Underlying Social Behavior. Cell 157, 1535–1551 (2014).

3. Lerner, T. N. et al. Intact-Brain Analyses Reveal Distinct Information Carried by SNc Dopamine Subcircuits. Cell 162, 635–647 (2015).

4. Wang, Y., DeMarco, E. M., Witzel, L. S. & Keighron, J. D. A selected review of recent advances in the study of neuronal circuits using fiber photometry. Pharmacol. Biochem. Behav. 201, 173113 (2021).

5. Lu, L. et al. Wireless optoelectronic photometers for monitoring neuronal dynamics in the deep brain. Proc. Natl. Acad. Sci. 115, E1374–E1383 (2018).

6. Guo, Q. et al. Multi-channel fiber photometry for population neuronal activity recording. Biomed. Opt. Express 6, 3919–3931 (2015).

7. Kim, C. K. et al. Simultaneous fast measurement of circuit dynamics at multiple sites across the mammalian brain. Nat. Methods 13, 325–328 (2016).

8. Pisano, F. Depth-resolved fiber photometry with a single tapered optical fiber implant. Nat. Methods 16, 12 (2019).

9. Sych, Y., Chernysheva, M., Sumanovski, L. T. & Helmchen, F. High-density multi-fiber photometry for studying large-scale brain circuit dynamics. Nat. Methods 16, 553–560 (2019).

10. Bruno, C. A. et al. pMAT: An Open-Source, Modular Software Suite for the Analysis of Fiber Photometry Calcium Imaging. http://biorxiv.org/lookup/doi/10.1101/2020.08.23.263673 (2020) doi:10.1101/2020.08.23.263673.

11. Holly, E. N. et al. Striatal Low-Threshold Spiking Interneurons Regulate Goal-Directed Learning. Neuron 103, 92–101.e6 (2019).

12. Muir, J. et al. In Vivo Fiber Photometry Reveals Signature of Future Stress Susceptibility in Nucleus Accumbens. Neuropsychopharmacology 43, 255–263 (2018).

